# Psilocybin induces sex- and context-specific recruitment of the stress axis

**DOI:** 10.1101/2025.01.06.631556

**Authors:** Sarah Gibson Cook, Stephanie Lee, Emma Ference, Yuecheng Shi, Mijail Rojas-Carvajal, Rene Hen, Jaideep S. Bains, Tamás Füzesi, Gergely F. Turi

## Abstract

Psychedelics have reemerged as potential treatments for mental health disorders, yet their impact on stress-related brain regions remains poorly understood. Here, we provide the first real-time, *in vivo* evidence of psilocybin-induced neuronal activation, specifically in hypothalamic corticotropin-releasing hormone neurons. Notably, psilocybin elicited more pronounced responses in female mice and produced context-related alterations in threat assessment. Our findings provide valuable insight into the impact of psychedelics on a key stress center in the brain.

## MAIN

Psychedelics have been used therapeutically for millennia^1^, and recent clinical trials have shown promise in their ability to treat stress-related mental health disorders, including depression and post-traumatic stress disorder (PTSD)^2,3^. These conditions involve dysregulation of the hypothalamic-pituitary-adrenal (HPA) axis and its associated stress hormones, particularly cortisol (corticosterone in rodents [CORT])^4,5^. Although it is established that psychedelics induce stress hormone secretion^6–8^, their impact on the primary regulators of CORT, corticotropin-releasing hormone neurons in the paraventricular nucleus of the hypothalamus (CRH^PVN^)^9,10^, is largely unknown.

The multifaceted nature of psychedelic responses reveals the variability of emerging treatments. Though stress-related disorders show considerable sex differences, with women experiencing higher rates of both depression and PTSD^11,12^, psychedelic clinical trials have not yet reported sex-specific effects. Conversely, mouse studies using psychedelics have found significant sex-related variations, particularly in hallucinogenic potential^13,14^ and effects on threat responses^15,16^. Additionally, the efficacy of psychedelic-assisted therapy can be heavily influenced by the individual’s mindset and the environmental setting of the experience, a phenomenon known as “set and setting” ^17,18^. Still, translating these complex psychological and contextual factors to animal models has proven challenging.

This study explores the effects of psilocybin (PCB) on CRH^PVN^ neurons using *in situ* hybridization and single-fiber photometry in mice. We validate that CRH^PVN^ neurons express serotonergic 5-HT2A receptors (5HT2Ars) and provide the first evidence of *in vivo*, real-time neuronal activation following PCB administration. We report that female animals show more substantial behavioral, endocrine, and neuronal responses to PCB, and that PCB-induced CRH^PVN^ activation is sensitive to contextual changes. Our work emphasizes the need for comprehensive *in vivo* investigation to better understand the nature and diversity of psychedelic experiences.

We first probed for behavioral changes in response to PCB to confirm the psychoactive effects of the drug. Wild-type mice received a single IP injection of vehicle or PCB (3 mg/kg). Head-twitch response (HTR), a potential proxy for hallucinogenic effects^13,14^, along with time spent rearing and grooming were monitored for 30 minutes post-injection. Consistent with previous research^8^, we observed an increase in HTR frequency in the PCB-treated group compared to vehicle controls, indicating a strong hallucinogenic reaction at this dose. Peak HTR occurred between 5 to 10 minutes after injection and returned to near baseline levels within 25 to 30 minutes (Fig. 1a). Comparison between sexes revealed a significantly higher HTR in females (Fig. 1b). We also noted distinct behavioral differences between vehicle and PCB-treated mice, including reduced time spent rearing and grooming after PCB (Fig. 1c,d).

**Fig. 1.**
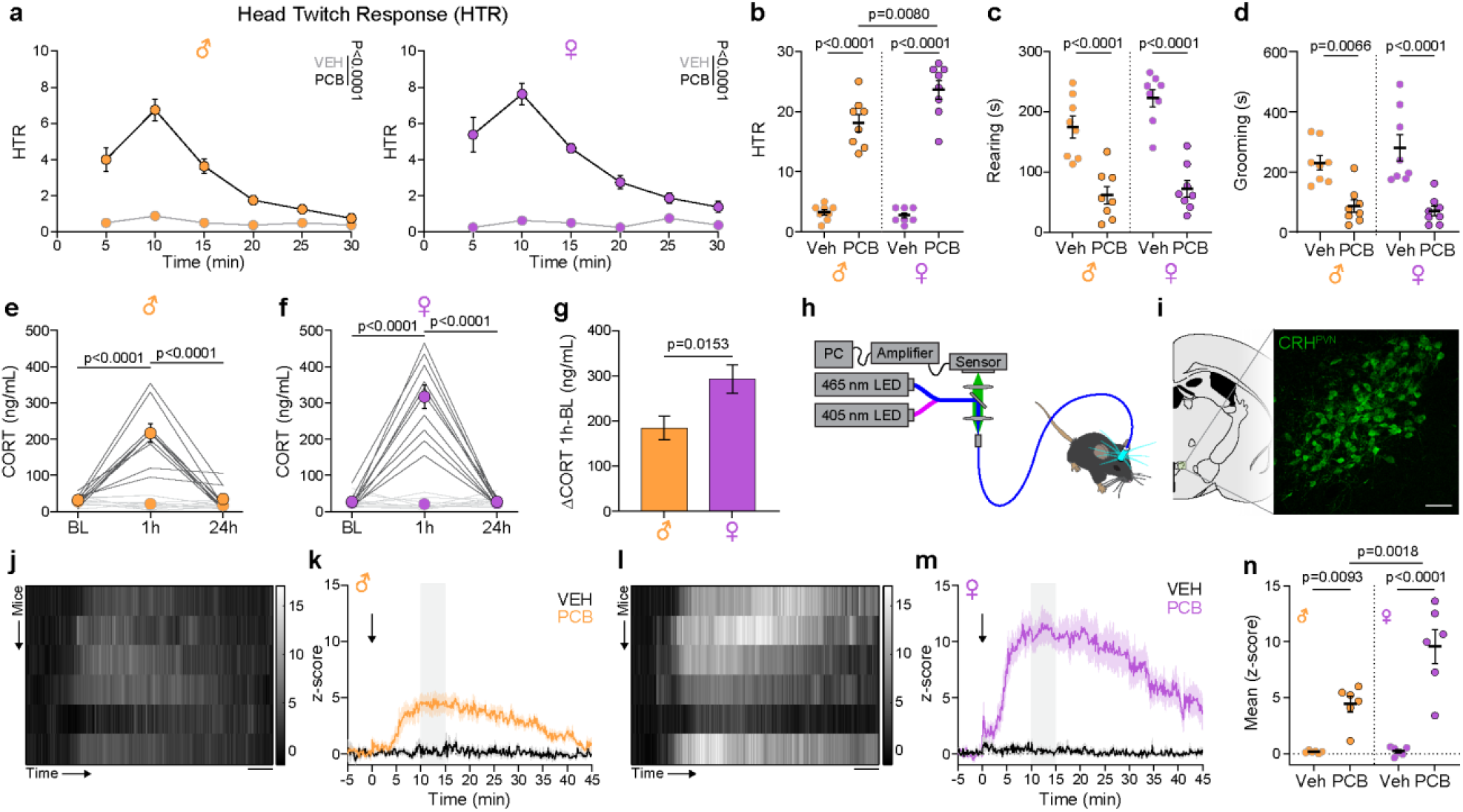
PCB induces behavioral, endocrine, and neuronal changes in a sex-specific manner. **a**, Timeline of head-twitch response (HTR) following vehicle or PCB (3 mg/kg, i.p.) male (orange) and female (purple) mice (Males: 5 min: p < 0.0001, 10 min: p < 00001, 15 min: p < 0.0001, 20 min: p = 0.0301, 25 min: p = 0.7185; 30 min: p > 0.9999, time x column factor: p > 0.0001, F [5, 70] = 22.07, n = 8 mice per group, Two-way ANOVA, Tukey’s post-hoc test); (Females: 5 min: p = 0.0053, 10 min: p < 00001, 15 min: p < 0.0001, 20 min: p = 0.0007, 25 min: p = 0.0403, 30 min: p = 0.1261, time x column factor: p > 0.0001, F [5, 70] = 20.34, n = 8 mice per group, Two-way ANOVA, Tukey’s post-hoc test). **b**, PCB-induced HTR is significantly higher in females (Males Veh vs. PCB: p < 0.0001, Females Veh vs PCB: p < 0.0001, PCB Males vs Females: p = 0.0080, F [3, 28] = 90.65, n = 8 mice per group, One-way ANOVA, Tukey’s post-hoc test); PCB also has significant effect on other stereotypic behaviours like (**c**) rearing (Males Veh vs. PCB: p < 0.0001, Females Veh vs PCB: p < 0.0001, F [3, 28] = 26.41, n = 8 mice per group, One-way ANOVA, Tukey’s post-hoc test); and (**d**) grooming (Males Veh vs. PCB: p = 0.0066, Females Veh vs PCB: p < 0.0001, F [3, 28] = 13.36, n = 8 mice per group, One-way ANOVA, Tukey’s post-hoc test). **e**, Plasma corticosterone concentrations were increased 1 hr following PCB and returned to baseline levels within 24 hrs (Males PCB BL vs 1 hr: p < 0.0001, PCB 1hr vs 24 hr: p < 0.0001, F [5, 54] = 44.33, n = 10 mice per group, One-way ANOVA, Tukey’s post-hoc test); (Females PCB BL vs 1 hr: p < 0.0001, PCB 1hr vs 24 hr: p < 0.0001, F [5, 54] = 70.34, n = 10 mice per group, One-way ANOVA, Tukey’s post-hoc test), and (**f**) this effect was stronger in female animals (p = 0.0153, t = 2.679, 10 mice per group, Unpaired, two-tailed t-test). **h**, Schematic depiction of single-fiber photometry recording of CRH^PVN^ neurons. **i**, Brain map and confocal image show the expression of GCaMP6f in CRH^PVN^ neurons within transgenic animals (scale bar = 50 µm). **j-m**, PCB injection induces a robust and enduring increase in the photometry signal recorded from CRH^PVN^. Heatmaps of CRH^PVN^ responses are shown for individual (**j**) male and (**l**) female animals (scale bar = 5 min). Grouped z-score values of photometry signals 5 mins before and 45 mins following vehicle or PCB injection. Comparisons show PCB induced CRH^PVN^ activation in both (**k**) male and (**m**) female mice (Males: p < 0.0001, t = 27.08, n = 6 mice per group, Unpaired, two-tailed t-test); (Females: p < 0.0001, t = 32.59, n = 6 mice per group, Unpaired, two-tailed t-test). **n**, Comparisons between sexes of mean z-score during the 10-15 min interval post-injection (**k**,**m**, grey shading) reveal significant differences (Males Veh vs. PCB: p = 0.0093, Females Veh vs. PCB: p < 0.0001, PCB male vs female: p = 0.0018, F [3, 20] = 27.88, n = 6 mice per group, One-way ANOVA, Tukey’s post-hoc test). Data are mean ± s.e.m.

To assess the impact of PCB on the stress axis, we collected peripheral blood samples and measured CORT levels before and after vehicle or PCB administration. PCB treatment induced a significant increase in CORT levels one-hour post-injection, which returned to baseline levels 24 hours later (Fig. 1e,f), confirming prior studies^6–8^. Further analysis stratified by sex revealed a significant difference within the PCB-treated group, with female mice exhibiting significantly higher CORT responses than males (Fig. 1g).

Because CORT secretion is driven by CRH^PVN^ neuron activation^9,10^, we next sought to monitor the activity of these neurons during the psychedelic experience. We used transgenic animals expressing GCaMP6f in CRH^PVN^ neurons and single-fiber photometry to track calcium changes as a proxy for population activity following vehicle or PCB treatment (Fig. 1h,I, see Extended Data Fig. 1). PCB elicited a robust and sustained calcium increase in CRH^PVN^ neurons, peaking at 10 - 12 minutes and returning to baseline after 45 minutes (Fig. 1j-m). Females showed significantly higher activity than males in the 10 to 15 min window following PCB injection, consistent with CORT measurements (Fig. 1n, see Fig. 1g).

A significant challenge in researching psychedelic compounds has been in identifying their specific neuronal targets. Psilocin, the active metabolite of PCB, shows strong affinity for 5-HT2Ars^19,20^ which are required for the HTR and hallucinogenic effects of psychedelics^19,21–23^. Furthermore, 5-HT2Ars within the PVN mediate neuroendocrine responses to the psychedelic and 5-HT2Ar agonist, DOI (1-(2,5-dimethoxy-4-iodophenyl)-2-aminopropane)^24^. Thus, we set out to determine the anatomical basis of a possible effect of PCB on CRH^PVN^ neurons, including sex-specificity. Neuronal tissue obtained from 5-HT2Ar-Cre mice crossed to the tdTomato Ai14 reporter line displayed red fluorescence signal in 5-HT2Ar expressing neurons in multiple brain regions (Fig. 2a). Specifically, tdTomato-expressing cells were observed in both the parvocellular and magnocellular divisions across the antero-posterior extent of the PVN (Fig. 2b). Additionally, extensive tdTomato expression was detected in the outer layer of the median eminence (Fig. 2c), where the axons of the parvicellular hypophysiotropic systems are located. Single RNAScope *in situ* hybridization using a CRH-specific probe confirmed overlap between tdTomato- and CRH-expressing cells (Fig. 2d).

**Fig. 2.**
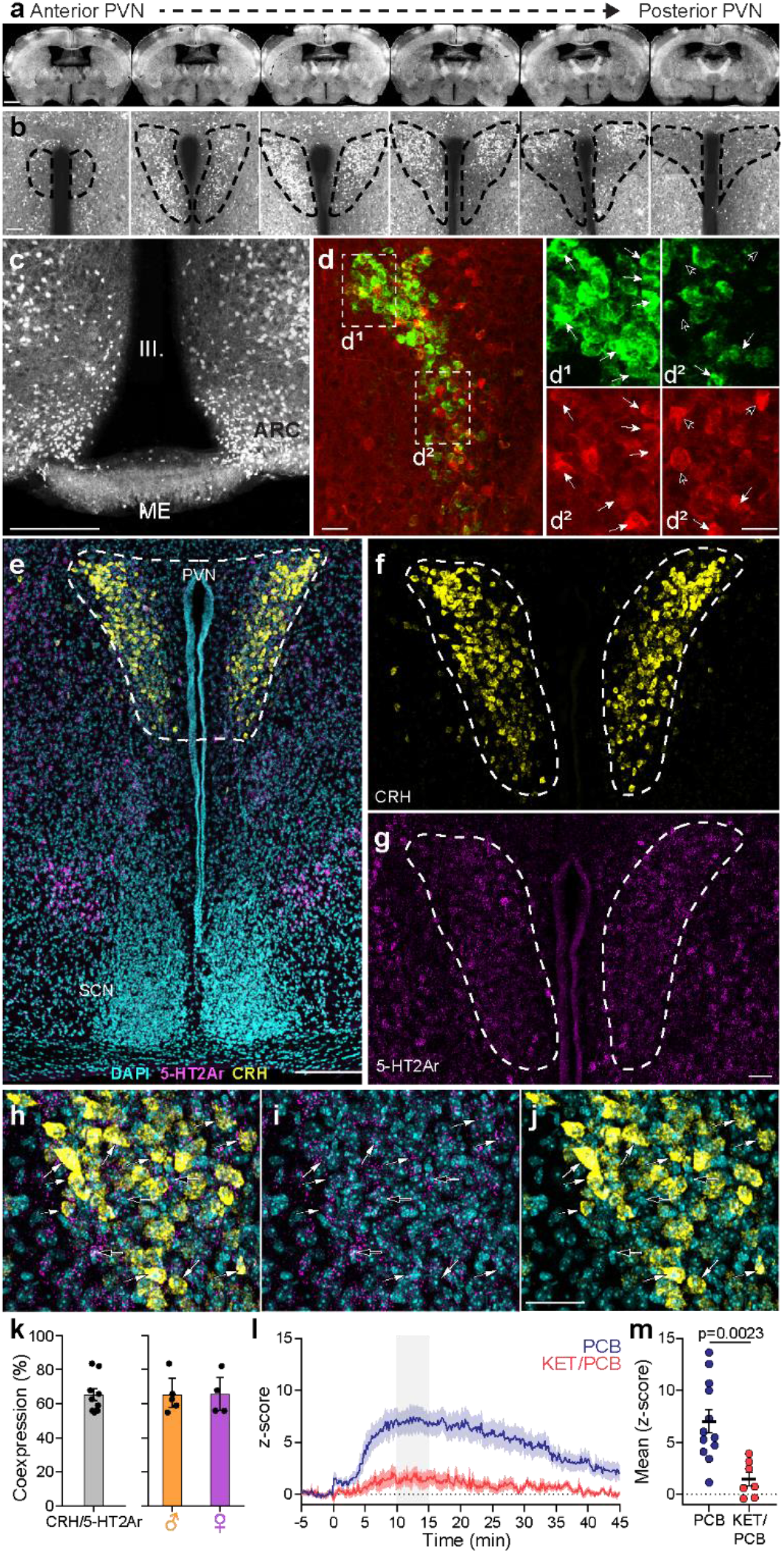
5-HT2Ar expression in the hypothalamus. **a-b**, Tissue sections from litters of 5-HT2Ar-Cre mouse crossed with Ai14D reporter line show expression of tdTomato in the antero-posterior extension of the PVN. **c**, tdTomato expression is also present in the outer layer of the ME where the axon collaterals of the hypophysiotropic systems are located. **d**, CRH in situ hybridization show overlap between CRH positive and tdTomato positive cells. (**d1**) and (**d2**) High magnification images correspond to areas delineated with white rectangles in (**d**). **e**, Representative image of PVN following detection of CRH (yellow), 5HT2Ar (magenta) and DAPI (cyan) RNA with dual label in situ hybridization. **f-g**, High magnification images representing the area in (**e**). **h-j**, Representative dual label in situ hybridization from the PVN. White arrows point to CRH/5-HT2Ar positive cells, empty arrows point to single labelled 5-HT2Ar+ cells. **k**, Quantification of dual labelled images revealed that 65.21±2.92% of the CRH neurons express 5-HT2Ar (n = 9 mice [5 males/4 females]). There is no statistical difference in the 5-HT2Ar between sexes. **l**, Pretreatment of CRH^PVN^ photometry mice with ketanserin (2 mg/kg) 1 hr before vehicle or PCB injection reduced PCB-induced CRH^PVN^ activation. **m**, Comparison between mean z-score in the 10-15 min post-injection time window (**l**, grey shading) shows significance differences between PCB and KET/PCB treated groups (p = 0.0023, t = 3.574, n = 12 mice [6 males/6 females] for PCB group, 6 mice [3 males/3 females] for KET/PCB group, Unpaired, two-tailed t-test). Scale bars: a – 1 mm; b – 150 µm; c – 250 µm; d – 50 µm; d1, d2 – 30 µm; e - 200µm; g,j - 50µm. PVN – paraventricular nucleus, SCN – suprachiasmatic nucleus, ME – median eminence, III. – third ventricle. Data are mean ± s.e.m.

As this breeding approach will also label cells which only transiently express 5-HT2Ars during development, we used dual label RNAscope hybridization to quantify the overlap between 5-HT2Ars and CRH. The signal distribution of the 5-HT2Ar RNA recapitulated the tdTomato pattern seen with the transgenic approach (Fig. 2e-g). Quantification of the dual-labeled sections revealed that 65% of the CRH^PVN^ neurons express 5-HT2Ar RNA (Fig. 2h-k). Notably, separating the sexes did not reveal differences in expression levels (Fig. 2k). To investigate the relevance of 5-HT2Ar expression in the PCB-triggered HPA cascade, we employed single-fiber photometry and found that pre-treating the mice with ketanserin (2 mg/kg), a 5-HT2Ar antagonist, drastically attenuated the activation of CRH^PVN^ neurons by PCB (Fig. 2l,m).

In humans, environmental setting is an important factor in psychedelic experiences^17,18^, however, this concept has not been extensively studied in animal models. Here, we explored whether the PCB-induced CRH^PVN^ activation is context-dependent by placing mice in a novel setting 10 minutes after vehicle or PCB (Fig. 3a). Both vehicle- and PCB-treated mice showed an increase in stereotypical exploratory behaviors including time spent walking and rearing when moved from their home-cage to the novel context (Fig. 3b,c), indicating that both groups can recognize the change in environment. Consistent with previous observations^25–27^, vehicle treated mice showed an increase in CRH^PVN^ activity during handling. Placement in the new context elicited an increase in activity that was sustained for the duration of the exposure (Figure 3d,e). In contrast, PCB-treated mice showed no increase in CRH^PVN^ activity during handling (Figure 3f) and a significant decrease in CRH^PVN^ activity in the novel setting (Figure 3g). Previous work has shown that CRH^PVN^ neurons demonstrate a sustained increase in activity when mice are placed in a novel, neutral context, likely as a means of threat anticipation^25^. Thus, the divergent responses in our results suggest that PCB affects the ability of CRH^PVN^ neurons to accurately interpret and respond to potential threat.

**Fig. 3.**
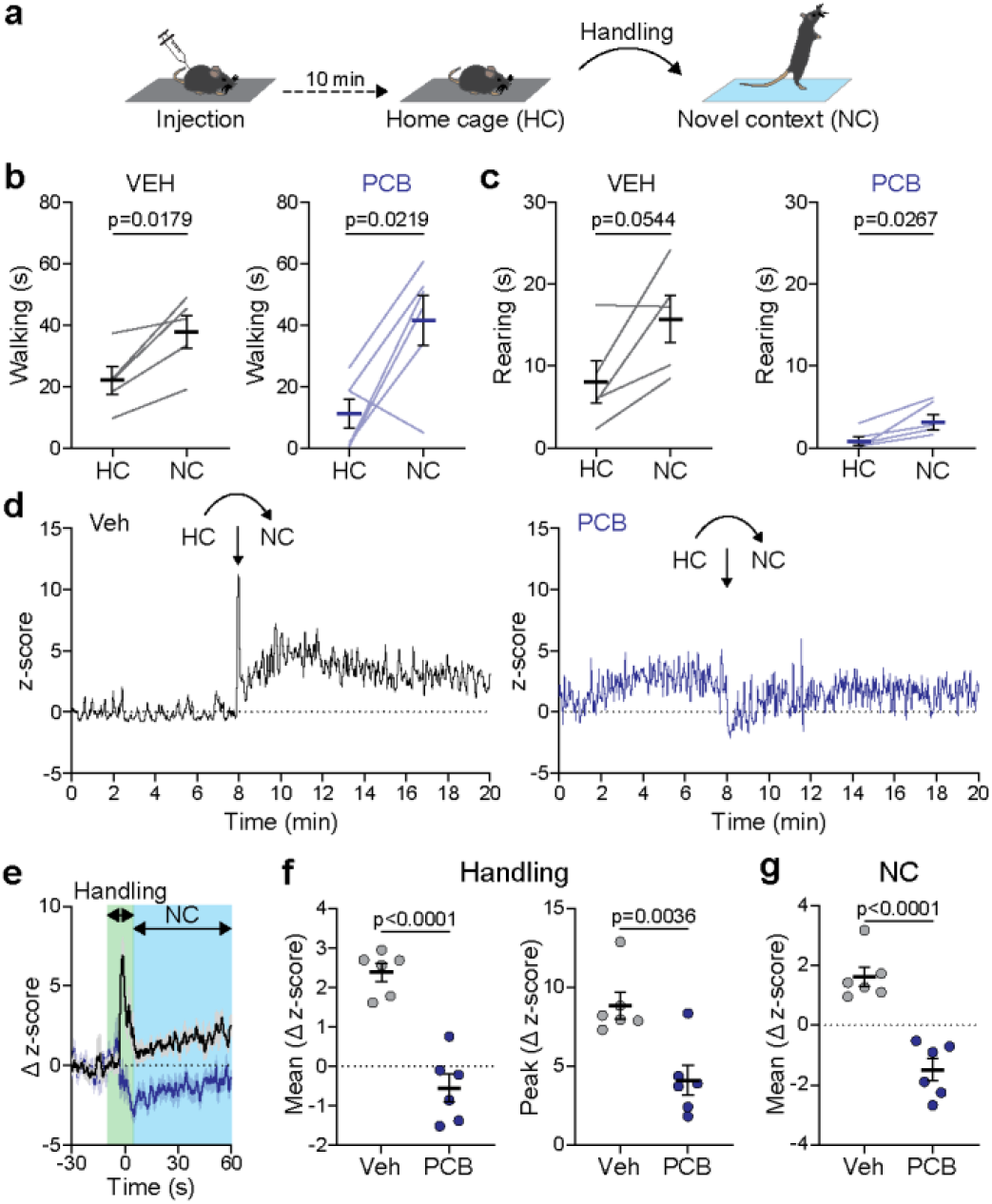
*In vivo* PCB-induced CRH^PVN^ activation is sensitive to contextual changes. **a**, Schematic illustration of context experiment: 10 minutes after vehicle or PCB injection in home-cage (HC), mice were moved from HC to novel context (NC). Time spent (**b**) walking and (**c**) rearing was quantified in the 3 min window before and after moving to the NC (Walking: VEH: p = 0.0179, t = 3.874, PCB: p = 0.0219, t = 3.282, n = 5-6 mice per group [2-3 males/3 females], Paired, two-tailed t-test); (Rearing: VEH: p = 0.0544, t = 2.695, PCB: p = 0.0267, t = 3.105, n = 5-6 mice per group [2-3 males/3 females], Paired, two-tailed t-test). **d**, Representative fiber photometry traces following vehicle or PCB treatment showing the change in CRH^PVN^ activity after moving from HC to NC. **e**, Changes in z-score for averaged photometry traces 30 sec prior to and 60 sec following moving from HC to NC used for further analysis of handling (green) and NC (blue) responses. Differences were seen comparing traces from vehicle and PCB-treated mice in (**f**) mean change in z-score and peak z-score for 15 second window during handling (Mean: p < 0.0001, t = 7.016, Peak: p = 0.0036, t =3.787, n = 6 mice per group [3 males/3 females], Unpaired, two-way t-test) and (**g**) mean change in z-score for 55 sec window following the move into NC (p < 0.0001, t = 6.282, n = 6 mice per group [3 males/3 females], Unpaired, two-way t-test). Data are mean ± s.e.m.

Our findings demonstrate that PCB activates the HPA axis through recruitment of CRH neurons in the PVN, leading to elevated CORT levels for up to an hour after administration. This robust effect is more pronounced in female mice, aligning with known sex differences in stress-related endocrine responses^28^. The PCB-dependent activation of CRH^PVN^ neurons relies on 5HT2Ars, widely considered the principal mediator of psychedelics’ hallucinogenic effects^21–23^. This connection suggests that the psychoactive properties of PCB may contribute to the activation of the stress response.

Sex differences emerged as a significant factor in our research, with female mice exhibiting stronger behavioral, endocrine, and neurological reactions to PCB. This sex-based disparity, often overlooked in clinical studies, has particular relevance for understanding psychedelics’ effects on brain function and behavior. The pronounced sex-specific responses suggest the need for sex-stratified clinical trial protocols, potentially tailoring dosages based on individual physiological stress responses. Our findings underscore the importance of investigating how sex influences both the stress and serotonergic systems, as these interactions may shape subjective psychedelic experiences and could inform more personalized therapeutic approaches.

Though our work reveals an association between the serotonin system and the HPA axis in response to PCB, some questions persist. We have confirmed 5HT2Ars are expressed in CRH^PVN^ neurons with anatomical tools, however, the precise mechanism remains elusive. Additional electrophysiological and molecular studies could examine whether this effect is direct or involves intermediate pathways. Furthermore, investigating how PCB alters the neural circuits that typically link CRH^PVN^ activity to stress-related behaviors could provide insights into how psychedelics modify established behavioral patterns. Also, exploration into whether acute HPA axis activation contributes to the efficacy of psychedelics may illuminate mechanisms that could transform stress responses into therapeutic opportunities.

CRH^PVN^ neurons typically show an increase in activity in response to adverse or novel stimuli^25–27^. Our findings reveal a surprising reduction in activity in a new setting when PCB is administered, suggesting a fundamental shift in established stress response mechanisms. This altered discernment of threat may be a key mechanism underlying psychedelics’ potential to mitigate stress consequences. Additionally, the observed flexibility aligns with emerging research on psychedelics’ capacity to disrupt fixed brain patterns and promote cognitive and neurological adaptability^29,30^. By inducing increased flexibility within the stress axis, psychedelics may offer a novel therapeutic approach for reshaping maladaptive stress responses.

This study represents one of the first demonstrations of real-time, psychedelic-induced setting effects in an animal model, providing insight into a concept long discussed in human research but rarely examined at the pre-clinical level. The suppression of PCB-induced CRH^PVN^ activation by a simple setting change illustrates how markedly environmental setting can shape psychedelic responses, highlighting the importance of treatment setting design in psychedelic-assisted therapy. Our findings suggest that the therapeutic effects of psychedelics extend beyond molecular interactions to involve sex-related and contextual factors, calling for more advanced *in vivo* tools to capture their complexity.

In conclusion, our research provides meaningful insight into the neurobiological effects of PCB on the stress system. The unexpected dissociation between CRH^PVN^ activation and typical stress-related responses, coupled with the context-dependent effects, suggests that PCB may create a unique state of HPA axis flexibility. This observation could be critical to the therapeutic potential of psychedelics, allowing for the reorganization of rigid patterns of thought and behavior. Our work positions psychedelics not merely as pharmacological treatments, but as potential tools for recalibration of stress-related brain circuits. By exploring further stress response modulation by psychedelics, we could reveal novel interventions for stress-related disorders and leverage the adaptability of our neuroendocrine system.

**Extended Data Figure. 1.**
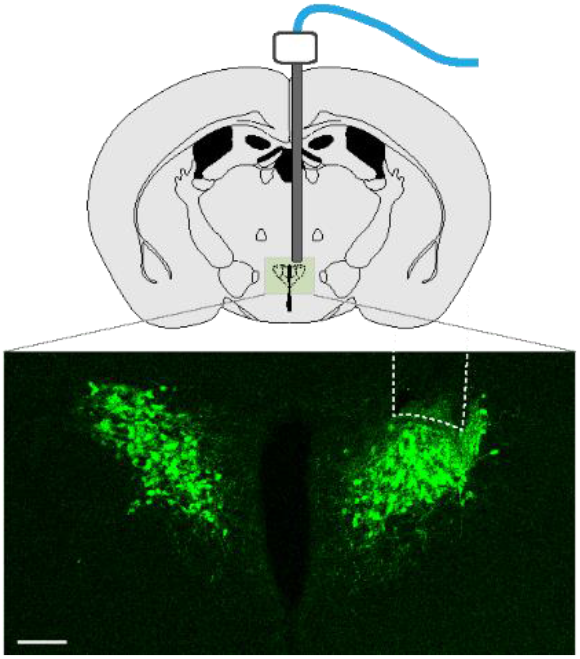
Implantation for fiber photometry. Schematic illustration of ferrule placement and low magnification confocal image showing the fiber photometry ferrule implantation site (dashed line) dorsal of the paraventricular nucleus of the hypothalamus (PVN, green). Scale bar = 100 µm.

## METHODS

### Mice

All procedures were carried out in accordance with the US National Institute of Health (NIH) guidelines for the care and use of laboratory animals and approved by the Animal Care and Use Committees of New York State Psychiatric Institute and the University of Calgary Animal Care and Use Committee. For fiber photometry and behavioral tracking experiments the offsprings of *Crh-IRES-Cre* (*B6(Cg)-Crhtm1(cre)Zjh/J*; stock number 012704) and *Ai148 (Ai148(TIT2L-GC6f-ICL-tTA2)-D*; stock number 030328) animals were used. Mice were obtained from Jackson Laboratories. For anatomy experiments the following lines were used: *Cg-Gt(ROSA)26Sor*^*tm14(CAG-tdTomato)Hze*^*/J* (stock number 007914) crossed with 5-HT2a-Cre (B6.Tg(Htr2a-Cre)KM208Gsat, MGI ID 5435493), and ArcCreERT2 x R26R-STOP-floxed-EYFP^2^ for RNAscope.

For behavioral experiments, mice were group-housed until the first experiment, then single-housed for at least 7 days on a 12:12 h light/dark schedule (lights on at 07:00 hours) with ad libitum access to food and water. All subjects were randomly assigned to different experimental conditions used in this study. For behavioral experiments, mice were 8-12 weeks old. For fiber photometry, mice were 6-8 weeks old at the time of surgery. All efforts were made to minimize animal suffering and the number of animals used.

### Materials and Drugs

Psilocybin (PCB) was purchased from Toronto Research Chemicals, dissolved in DMSO (100 mM), aliquoted and stored in a -80° freezer. On the morning of each experiment, vehicle (1% DMSO in 0.9% saline) and PCB (diluted to 3 mg/kg in 0.9% saline) were prepared and administered using intraperitoneal (i.p.) injections at 2.5 µL/g body weight. The 3 mg/kg dose was chosen as it has been shown to induce acute CORT release in male mice^8^. Ketanserin was purchased from Sigma-Aldrich and prepared and administered in a similar manner to PCB except diluted to 2 mg/kg, a dose sufficient to reduce PCB-induced head-twitch responses^31^.

### Behavioral recordings and analysis

Head twitch response (HTR; a rapid, involuntary movement of the mouse’s head with little or no involvement of the trunk) is widely utilized as a behavioral proxy in rodents for human hallucinogenic effects and can reliably differentiate between hallucinogenic and non-hallucinogenic 5-HT2Ar agonists^13^. Before recordings, mice were exposed to the testing chamber for at least 10 mins, 1 day prior to testing to minimize novelty effects. The testing chamber was cleaned with a 70% ethanol solution between experiments and bedding was changed to eliminate odor from other mice. Mice were video monitored for 30 minutes directly after vehicle or PCB injection using a top-mounted infrared light and camera in a plexiglass testing chamber (25.5 × 12.5 × 12.5 cm [L x W x H], with 5 cm of fresh bedding) to record behavior and returned to their home-cage. Behavioral analysis to quantify HTR and time spend rearing and grooming was conducted by an individual blinded to subject treatment group using Behavioral Observation Research Interactive Software (BORIS, version 7, DOI: 10.1111/2041-210X.12584). Behavior during 5-minute time windows was quantified and plotted, i.e. 5 min = 0-5 minute interval.

### Corticosterone measurement

24 hrs before, 1 hr after, and 24 hrs after either vehicle or PCB administration in home-cage, blood from the tail vein was collected into ice-cold EDTA capillary tubes (Sarstedt) and centrifuged (Eppendorf centrifuge 5430 R, 8100 g, 4 °C, 20 min). Aliquots of plasma were stored at −20 °C until assayed using a DetectX Corticosterone Immunoassay kit (Arbor Assay). Plasma samples were run in triplicates on the same day.

### *In situ* hybridization

The mice (n=9; 5 males, 4 females) were transcardially perfused with 10 ml 0.1 mM phosphate buffered saline (PBS, pH=7.5) followed by 40 ml 4% paraformaldehyde (PFA) dissolved in PBS, then post-fixed overnight in 4% PFA. After fixation, the brains were sectioned using the vibratome at 50 μm thick coronal slices and stored in cryoprotectant at -20°C until the *in situ* hybridization.

Free-floating vibratome sections were washed 3 times for 15 minutes in PBS at room temperature. Sections were treated with RNAscope hydrogen peroxide for 10 minutes, then washed 3 more times in PBS (0.1 mM, pH 7.4). Sections were mounted on silane-treated slides (Sigma-Aldrich, 41122601), dried for 30–60 minutes at -20°C, and heat treated at 60°C for 30 minutes. Slides were sequentially immersed in 50%, 70%, and 100% ethanol for 5 minutes each, followed by air drying. Slides were incubated in pre-heated 1x Target Retrieval Reagent (ACD, 323110) at 99°C for 15 minutes, washed briefly in distilled water, and then immersed in 100% ethanol for 3 minutes. A hydrophobic barrier was drawn around tissues with an ImmEdge pen. Protease III (ACD, 322381) was applied for 30 minutes at 40°C. RNAscope probes were applied to the slides and incubated at 40°C for 2 hours, followed by two washes with 1x wash buffer (ACD, 310091). Slides were sequentially incubated with RNAscope AMP 1, AMP 2, and AMP 3, each at 40°C (AMP 1 and AMP 2 for 30 minutes, AMP 3 for 15 minutes), followed by washes with 1x wash buffer (ACD, 310091). HRP-C1 was added, and slides were incubated for 15 minutes at 40°C, then washed (ACD, 323110). TSA Vivid Fluorophore (520) was applied for 30 minutes at 40°C, followed by additional washes. A blocking solution was applied for 15 minutes. If a second probe was used, HRP-C2 was applied and followed by TSA Vivid Fluorophore (570 or 650) and blocking as with the C1 probe. DAPI was applied to the slides for 30 seconds, followed by mounting with a coverslip. Slides were dried in the dark for 30 minutes to overnight and stored at 2–8°C.

### Confocal imaging

Fluorescent confocal micrographs were captured using the Leica SP8 confocal microscope and Leica Application Suite X software. Predetermined settings optimized for the dyes were applied to scan all slices. Approximately four brain sections of each mouse were scanned at 10X magnification to identify PVN. A single frame Z-stack was taken from a representative region within each PVN at 20X magnification, with a step size ranging from 0.507 μm to 0.760 μm. To improve signal-to-noise ratio and obtain a clearer image, a frame average of 2 was applied during scanning. Z-stacks were captured from top to bottom of each slice due to slight variations in section thickness, although all brains were sliced into 50 μm sections using a vibratome.

### Quantification of *in situ* hybridization

Maximum projected images were used to analyze cell counts and expressed as the number of fluorescent cells/volume (μm^3^), with consideration given to the thickness of the Z stack. Throughout the experiment, the investigator remained blind to the treatment status. To manually analyze and count cells in images, the Fiji/ImageJ Cell Counter tool was used. If the images were too dark, we adjusted the image contrast to facilitate manual cell counting. To obtain the cells/volume (μm^3^) for analysis, the cell count was divided by the volume of the drawn ROI around the perimeter of the PVN. Channels were separated and signals in each were marked and counted. Channels were then remerged and cells with both signals, representing events of colocalization, were counted.

### Optical fiber implantation

For fiber photometry experiments, a 400 µm diameter mono fiber optic cannula (Doric Lenses, MFC_400/430/0.48_5mm_MF2.5_FLT) was implanted in *Crh-IRES-Cre*;*Ai148* mice, as previously described^25,26^. 6-8 weeks old mice were maintained under isoflurane anesthesia in the stereotaxic apparatus. Implantations were targeted dorsal to the PVN (AP, −0.7 mm; L, -0.2 mm from the bregma; DV, −4.2 mm from the dura) and were affixed to the skull with METABOND^®^ and dental cement. Experiments started after at least one week of recovery.

### Fiber photometry recording

Fiber photometry was used to record calcium transients from CRH neurons in the PVN of freely moving mice. After the recovery period, animals were handled for 5 minutes a day for 3 successive days and then habituated to the optic fiber in their home-cage (15 minutes a day) for 3 additional days. We recorded 20 minutes of baseline CRH^PVN^ neuron activity in the home-cage immediately before each drug administration for better bleaching correction and the first 5 minutes of recording were excluded from further analysis.

Doric fiber photometry system: Consisting of two excitation LEDs (465 nm and 405 nm from Doric) controlled by a LED driver and console, running Doric Studio software (Doric Lenses). The LEDs were modulated at 208.616 Hz (465 nm) and 572.205 Hz (405 nm) and the resulting signal demodulated using lock-in amplification. Both LEDs were connected to a Doric Mini Cube filter set (FMC5_E1(405)_E2(460-490)_F1(500-550)_S) and the excitation light was directed to the animal via a mono fiber optic patch cord (DORIC MFP_400/460/900-0.48_2m_FC/MF2.5). The power of the LEDs was adjusted to be 30 µW at the end of the patch cord. The resulting signal was detected with a photoreceiver (Newport model 2151).

### Fiber photometry data analysis

Fluorescent signal data was acquired at a sampling rate at around 100 Hz (Doric system). Data was then exported to MATLAB (Mathworks) for offline analysis using custom-written scripts. Briefly the 465/470 nm and 405 nm data were first individually fit with a second order curve which was then subtracted to remove any artifacts due to bleaching. Next a least-squares linear fit was applied to the 405 nm to align it with the 470 nm channel and then the change in fluorescence (ΔF) was calculated by subtracting the 405 nm Ca^2+^ independent baseline signal from the 470 nm Ca^2+^ dependent signal at each time point.

To minimize the impact of the handing and the upstate on the bleaching correction, the baseline and a segment from more than 40 min after injections were used for curve fitting. Additionally, the 30 second window immediately surrounding injection was subtracted from traces prior to analysis. For analysis, z-score calculation was performed using the following equation z=(F−F_0_)/σF. Where F is the test signal, F_0_ and σF are the mean and standard deviation of the baseline signal, respectively.

### Statistics

GraphPad Prism 10.0 software was used for statistical analysis. When comparing means from two dependent groups, different time points, paired t-test (two-tailed) were used. When comparing the means of two independent groups, unpaired t-test (two-tailed) were used. When comparing the means of multiple groups, parametric repeated measures, mixed effects model, one-way or two-way ANOVA were used followed by Tukey’s post-hoc test for multiple comparisons. Results are expressed as mean ± standard error of the mean, unless otherwise noted.

## DATA AVAILABILITY

The raw data that support the findings of this study are available from the corresponding author upon reasonable request.

## CODE AVAILABILITY

Scripts used to analyze fiber photometry is deposited here: https://github.com/leomol/FPA.

## ACKNOWLEDGEMENTS

We thank Mrs. Cheryl Breiteneder, Dr. Dinara Baimoukhametova, Ms. Alexis Passmore, and Mr. Rodney Barasi for expert technical support. We also thank Dr. Eleanor Simpson and Dr. Matthew Hill for access to materials. We are grateful for the support of the Cumming School of Medicine Optogenetics Core Facility.

This work was supported by an operating grant to J.S.B. from the Canadian Institutes for Health Research (FDN-148440) and the Brain Canada Neurophotonics Platform. M.R.C. was supported by the Izaak Walton Killam Memorial Doctoral Scholarship. G.F.T. and R.H. were supported by the Hope for Depression Research Foundation.

## AUTHOR CONTRIBUTIONS

S.G.C. designed and conducted the behavioral, endocrine, and photometry experiments, analysed data, and wrote the paper. M.J.C. contributed to the photometry experiments. S.L., E.F., Y. S. conducted the *in situ* hybridization experiments. R.H. and J.S.B. discussed the results and supervised the project. T.F. and G.F.T. designed experiments, analysed data, and contributed to the paper. Controlled drug license was acquired and *in vivo* behavioral and fiber photometry experiments were performed in the laboratory of J.S.B. Neuroanatomical work was conducted in the lab of G.F.T.

